# Pairs and Pairix: a file format and a tool for efficient storage and retrieval for Hi-C read pairs

**DOI:** 10.1101/2021.08.24.457552

**Authors:** Soohyun Lee, Carl Vitzthum, Burak H. Alver, Peter J. Park

## Abstract

**Summary:** As the amount of three-dimensional chromosomal interaction data continues to increase, storing and accessing such data efficiently becomes paramount. We introduce Pairs, a block-compressed text file format for storing paired genomic coordinates from Hi-C data, and Pairix, an open-source C application to index and query Pairs files. Pairix (also available in Python and R) extends the functionalities of Tabix to paired coordinates data. We have also developed PairsQC, a collapsible HTML quality control report generator for Pairs files.

**Availability:** The format specification and source code are available at https://github.com/4dn-dcic/pairix, https://github.com/4dn-dcic/Rpairix and https://github.com/4dn-dcic/pairsqc.

## 1 Introduction

A Hi-C assay combines chromosome conformation capture with high-throughput sequencing to interrogate genome-wide chromosome organization at high resolution. Modern Hi-C experiments (Rao, S.S.P. et al., 2014, Krietenstein et. al. 2020) can yield billions of paired genomic coordinates, representing a comprehensive set of three-dimensional DNA contacts in a cell population. This paired coordinate information can be utilized to explore interactions between specific genomic regions. We sought a file format optimal for storing and querying this information.

One of the most widely used options for storing pairs of read-level coordinates is BAM (Li, H. et al., 2009), a binary format that allows random access queries of genomic regions through Samtools. However, the BAM format does not support efficient querying of both coordinates in a pair, as it requires scanning the entire query result for the first coordinate of the pair, where the query result can span the whole genome. It also has an intrinsic redundancy of storing every pair of coordinates twice, so that the two coordinates are treated symmetrically. Moreover, BAM specifications are tightly defined and difficult to modify to circumvent these problems. A format used by Juicer (Durand, N.C. et al., 2016) called *merged_nodups.txt* stores pairs of genomic coordinates non-redundantly, but it is intended to be an intermediate format and is not optimized for storage or querying. Tabix (Li, H., 2011) is a software tool that offers similar indexing and querying functionality to Samtools on block-compressed tab-delimited text files, thus allowing for flexibility in the file format. However, similar to the BAM format, indexing or querying a second coordinate in a pair is not supported by Tabix. BEDPE (Quinlan, A.R and Hall, I.M., 2010) is a format for storing paired genomic regions, but it is not economical for storing massive read-level Hi-C data due to its constraints of required and reserved fields.

To overcome these issues, we designed the Pairs file format, which avoids redundancy in storing paired data, and Pairix, an extension of Tabix to index and query over pairs of genomic coordinates. We named this format ‘Pairs’ given the colloquial language that refers to this type of files as ‘pairs’ files.

## 2 Results

### 3.1 Pairs format

The Pairs format consists of header lines and data lines. Each data line corresponds to a pair of genomic coordinates. There are four required fields (chr1, pos1, chr2, and pos2) corresponding to two chromosomes and positions. To make it Pairix-compatible, the content should be sorted by chr1, chr2, pos1, then pos2. There are three reserved fields (readID and strand1/2, but could be left blank) and two reserved field names that can be used for optional fields. To reduce redundancy, the Pairs format stores Hi-C data in an upper-triangular matrix instead of the whole matrix, ensuring that the second coordinate of the pair is larger than the first one. If the data matrix is asymmetric, e.g., MARGI for RNA-DNA interactions (Sridhar, B. et al., 2017), the whole matrix could be stored.

A more detailed specification can be found at https://github.com/4dn-dcic/pairix/blob/master/pairs_format_specification.md. A Pairs file has the .*pairs.gz* extension after block-compression using bgzip (Li, H., 2011), and the extension of its Pairix index has .*pairs.gz.px2* (*px2* instead of *tbi*, a typical extension of a Tabix index).

### 3.2 Queries

Figure 1 summarizes examples of three types of queries; intra-chromosomal (e.g., getting all interactions between chrX:40,000,000-60,000,000 and chrX:80,000,000-100,000,000), inter-chromosomal (e.g., getting all interactions between chr1:100,000,000-300,000,000 and chrY:7,000,000-8,000,000) and 4C-like queries (e.g., getting all interactions between chr19:20,000,000-30,000,000 and any other region in the genome).

**Fig. 1.**
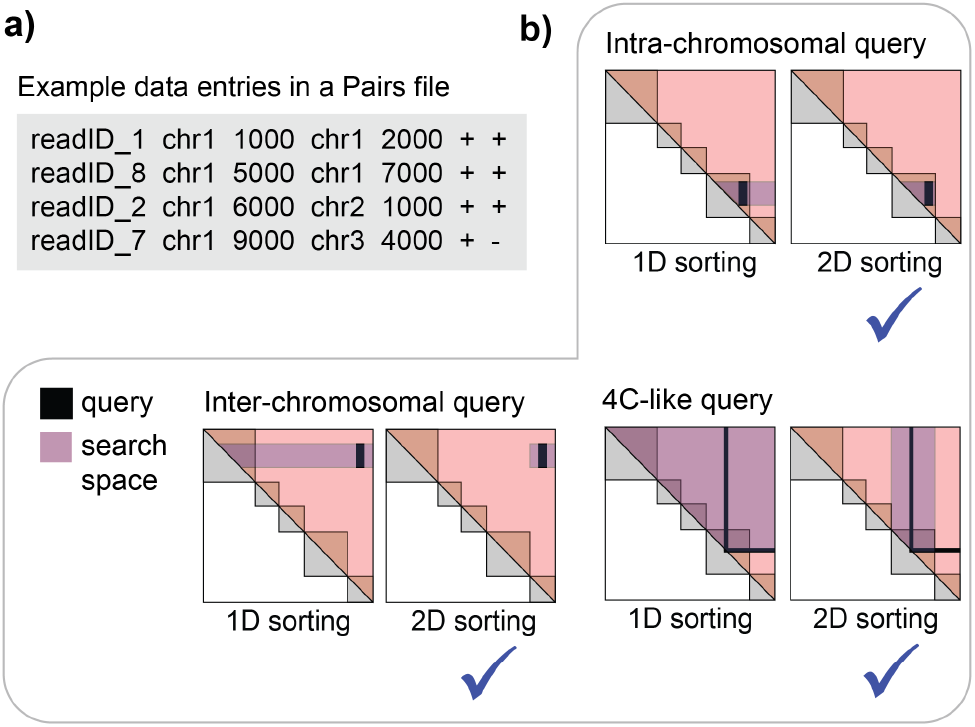
Pairs format and common types of queries for Pairs. **a) Pairs format, b)** comparison of search space between conventional 1-D sorting and the 2-D sorting adopted by Pairs and Pairix, for three types of queries over pairs of genomic loci. In all three scenarios, using a 2-D sorted file involves a smaller search space.

The figure illustrates the search spaces for each example query with the conventional ‘1D sorting’ (sorting by chr1 then pos1) and with the ‘2D sorting’ that we adopted for Pairs and Pairix (sorting by chr1, chr2, pos1 then pos2). Clearly, the 2D sorting is more efficient.

### 3.3 Pairix

Pairix has four main additional features compared to Tabix. First, it can use pairs of chromosomes as hash keys, depending on the sorting mechanism. Second, it can parse both single- and paired-region queries. Third, it accepts Pairs as the default format. Fourth, the number of lines in a file is stored in the index and can be retrieved instantly. Most of the original Tabix functionalities are preserved. However, due to the modifications in the index and query structures, Tabix and Pairix are not mutually interchangeable, i.e., Tabix cannot be used with a Pairix index or vice versa. Pairix as well as Pypairix and Rpairix (Python and R libraries) are available at https://github.com/4dn-dcic/pairix and https://github.com/4dn-dcic/Rpairix.

### 3.4 Performance of Pairix

Creating an index file is a one-time task and its run time increases linearly with data size, taking 0.68 seconds per million lines of a Pairs file plus 9 seconds of overhead on an AMD EPYC 7571 CPU. Memory requirement saturates at about 200 Mb.

The performance improvement in queries using random access compared to not using random access is substantial, as expected. For a minimal Pairs file that contains over 300 million lines, retrieving an entry takes ~1 second with Pairix, whereas scanning the whole file takes up to 30 minutes, depending on the genomic regions of interest.

In terms of the size, a 4DN Hi-C BAM file containing 3,285,241 reads is 870.55 MB, whereas the Pairs file containing 81.6% of the reads (2,679,253 reads) is only 38.08 MB.

### 3.5 PairsQC

PairsQC is an open-source quality control (QC) HTML report generator for Hi-C Pairs files based on Nozzle (Gehlenborg, N. et al., 2013) and D3 (Bostock, M. et al., 2011). The collapsible HTML report generated contains a summary table of various quality metrics and several QC plots for Hi-C. The source code and example reports can be found at https://github.com/4dn-dcic/pairsqc.

## Discussion

Pairs, Pairix and PairsQC have been used to process and store filtered read-level Hi-C and Hi-C-like data for more than 600 experiment sets for the 4D Nucleome consortium (Dekker, J. et al., 2017), with the results available at https://data.4dnucleome.org. The Pairs format is supported by Juicer (Durand, N.C. et al., 2016), Cooler (Abdennur, N. and Mirny, L.A., 2019) and Pairtools (https://github.com/mirnylab/pairtools).

## Acknowledgements

We thank Nezar Abdenur, Anton Goloborodko and other 4D Nucleome members for the valuable discussion. We also thank Scott Kallgren for incorporating GenomicRanges (Lawrence M, et al. (2013)) in Rpairix

